# MycorrhizaTracer: A BIOINFORMATIC PIPELINE FOR FUNGI AND PLANT CLASSIFICATION OF SANGER DNA SEQUENCES

**DOI:** 10.64898/2026.04.23.720352

**Authors:** T. D. Brekke, T. Weeks, R. A. Barber, I. Thomson, R. Gooda, R. Gargiulo, G. Delhaye, C. Andrew, J. Kowal, M. Bidartondo, L. M. Suz

## Abstract

Processing Sanger DNA sequences remains a routine yet technically demanding step in many biodiversity and ecological studies, particularly when barcoding large numbers of environmental samples. Manual inspection and editing of trace files, DNA sequence alignment, and classification using taxonomic reference databases is time-consuming, inconsistent, and prone to error. These challenges are compounded in studies involving degraded samples, in-house DNA sequencing, under-described taxa, or when investigators have limited access to computational tools.

We present MycorrhizaTracer, an open-source, fully automated pipeline for processing and taxonomically classifying large batches of Sanger sequencing chromatograms. We have optimized it for fungal and plant taxa, but it is adaptable across the tree of life. The pipeline performs quality trimming, consensus generation from bidirectional reads, taxonomic classification via BLAST, clustering, optional salvaging of low-quality sequences, and functional annotation of fungal taxa. Designed for scalability and ease of use, MycorrhizaTracer can process thousands of DNA chromatograms in a matter of hours without the need for an HPC. Accuracy and ecological relevance are ensured by features such as gene region-specific taxonomic filtering and sequence-based clustering of unclassified reads. By streamlining trace-to-taxon workflows, MycorrhizaTracer reduces the burden of manual curation, supports reproducibility, and enables efficient recovery of biodiversity data from Sanger sequences - particularly in field-based or resource-limited research contexts.

## Introduction

In ecological sampling and biodiversity research, Sanger sequencing continues to serve as a dependable and accessible tool for species identification. Its proven accuracy, straightforward protocols, and moderate costs support its widespread use in DNA barcoding and targeted marker studies. While high-throughput sequencing now dominates large-scale ecological genomics, the enduring relevance of Sanger sequencing reflects its strength in applications that require precise, individual-level sequence data.

Sanger sequencing remains a widely applied methodology for high-accuracy sequencing of single-nucleotide variants and for environmental surveys, especially when newer HTS technologies are either inaccessible or not feasible to utilise, given the study design. For example, in cases with focused identification or detection of target individual taxa (van der Linde et al. 2018; Cale et al. 2021), or where project scope, accessibility to other technologies, and/or budget limit the need for deeper sequencing, Sanger sequencing offers an affordable, long-read, fast-turnaround, and accessible alternative to HTS. In particular, ecological, conservation, and taxonomic research can still rely on Sanger-generated DNA barcodes to build or expand curated reference databases (Abarenkov et al. 2023; Ratnasingham et al. 2024), document species occurrences (Sting et al. 2019; Cantonwine et al. 2022), and train new scientists (Horn et al. 2020). Many widely used species-identification databases, such as UNITE for fungi (Abarenkov et al. 2023) were built from Sanger sequences ∼600–1,000 base pairs (bp) in length. This makes them a poor fit for long-read sequences of 2,500 bp or more (e.g., from PacBio), where regions extending beyond the reference provide no additional information for identification (though they may be useful for phylogenetic reconstruction). Conversely, most high-throughput sequencing relies on Illumina technology, which produces reads of at most ∼550 bp (paired-end 300 bp with a 50 bp overlap). This constraint has made the shorter ITS2 (Internal Transcribed Spacer) fragment (100-400 bp in fungi; Schoch et al. 2012) a common target for Illumina-based studies (Schoch et al. 2012; Olds et al. 2023; Winand et al. 2025). However, the UNITE database generally includes the complete ITS region (ITS1, 5.8S, and ITS2), which can be up to ∼1,200 bp in some fungi (Heeger et al. 2019) and which Sanger sequencing can typically capture in its entirety. Thus, for species identification against databases such as UNITE, Sanger read lengths are ideally matched, providing higher taxonomic resolution than is typically possible with Illumina while also avoiding the pitfall of the long-read technology where numerous bases are sequenced that fall outside the range of the current databases (though new long-read based databases such as Eukaryome (Tedersoo et al. 2024) are beginning to address this).

To aid studies that utilise Sanger sequencing for large numbers of samples, we present MycorrhizaTracer (from *<u>mycorrhiza</u>* identification of chromatogram *<u>trace</u>* files), a bioinformatic pipeline specifically designed for the high-throughput processing and classification of Sanger sequencing chromatograms optimised for plant and fungi identification. MycorrhizaTracer supports scalable, automated handling of chromatogram trace files, quality filtering, alignment, consensus calling, taxonomic assignment, clustering, and functional annotation of each fungal sample using FUNGuild (Nguyen et al. 2016). This pipeline facilitates the integration of Sanger-derived data into advanced biodiversity, ecology, and functional analyses. By providing a standardized and reproducible framework, our approach enables researchers to unlock the full value of existing and newly generated Sanger DNA datasets.

In this manuscript we demonstrate the utility of MycorrhizaTracer using a case study of thousands of DNA sequence chromatograms obtained from individual ectomycorrhizas from diverse taxa collected from soil samples across England. We demonstrate that, rather than being superseded entirely, Sanger sequencing and HTS should be viewed as complementary tools in the molecular ecology toolkit.

### Project description

MycorrhizaTracer is a bioinformatic pipeline to automate the processing, quality control, and taxonomic classification of large amounts of Sanger sequence data, optimized for fungal and plant DNA barcode sequences from the ITS region of the nuclear ribosomal genes and/or the *rbcL* gene. This pipeline is structured in five main steps following pre-pipeline preparations: (1) consensus calling, (2) classification, (3) clustering, (4) salvaging, and (5) functional annotation (Fig 1).

**Figure 1:**
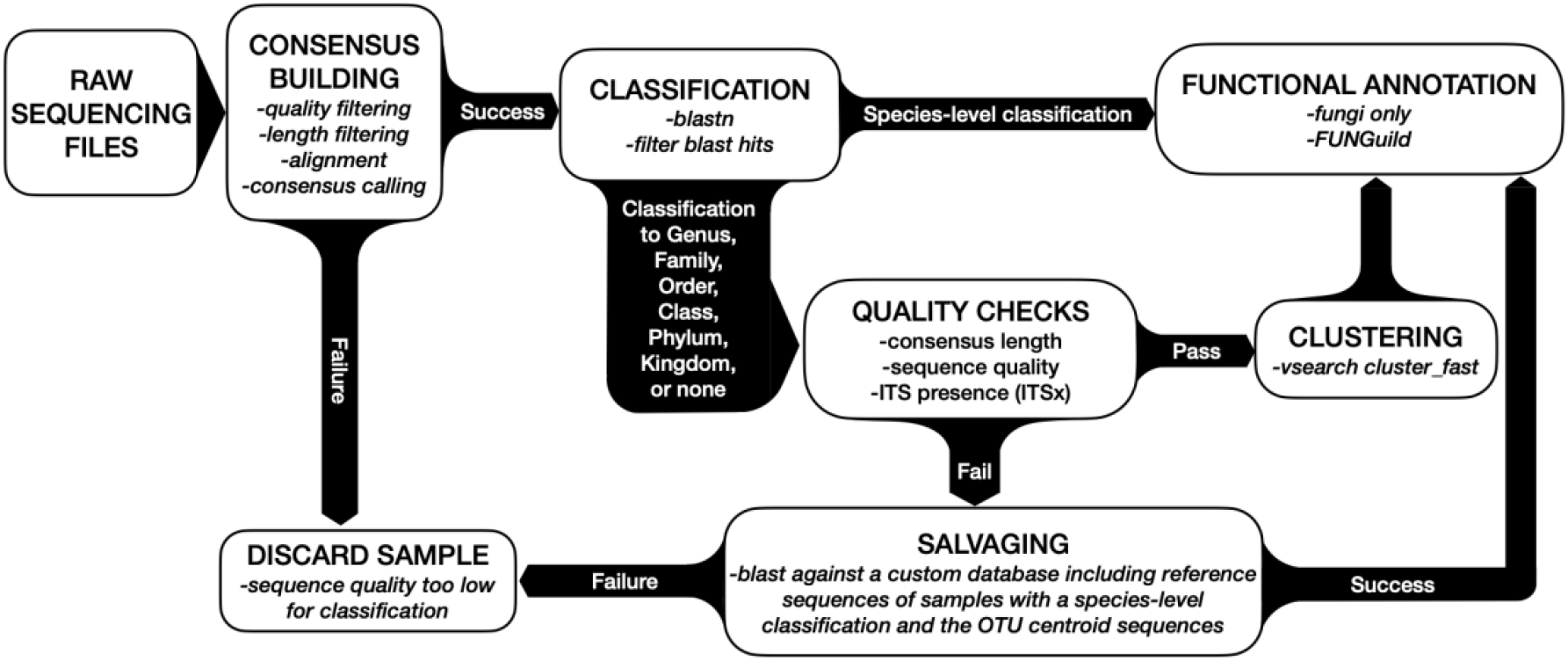
Pipeline step diagram. Each major component can be fine-tuned with command line options. RAW SEQUENCING FILES (chromatograms) are used for CONSENSUS BUILDING where they are filtered, aligned, and converted into a consensus based on either one of or both forward and reverse sequencing reads. Failure to generate a consensus, results in discarding the sample, as the sequence quality is too low. Consensus sequences are then passed to the CLASSIFICATION module where they are blasted against the specified database, and the blast hits are filtered to assign the sample a taxonomy. All samples that did not achieve a species-level taxonomic classification undergo a series of QUALITY CHECKS for length, sequence quality, and (if the target region includes ITS) the presence of the motifs flanking the ITS region. Samples passing these strict quality checks are sent for CLUSTERING into Operational Taxonomic Units (OTUs). Samples failing the quality checks are directed to SALVAGING where they are blasted against a custom database that includes both, the reference sequences linked to the samples with species-level taxonomies, and the cluster centroids of the OTUs. All fungal samples that either (1) have a species-level classification, (2) were clustered into OTUs, or (3) were successfully salvaged, can optionally undergo FUNCTIONAL ANNOTATION with *FUNGuild*.

#### Pre-pipeline

The pipeline by default expects that each sample has been sequenced in the forward and reverse directions for some or all of the following genomic regions: full ITS (fungi), ITS2 (plants), and *rbcL* (plants). We have tested specific primer combinations and databases (Table 1), but the pipeline can be adapted to any taxa and genomic region by using an appropriate database (i.e., specific to that DNA region/marker) and PCR primers that target the correct genomic region in the correct taxa.

**Table 1:**
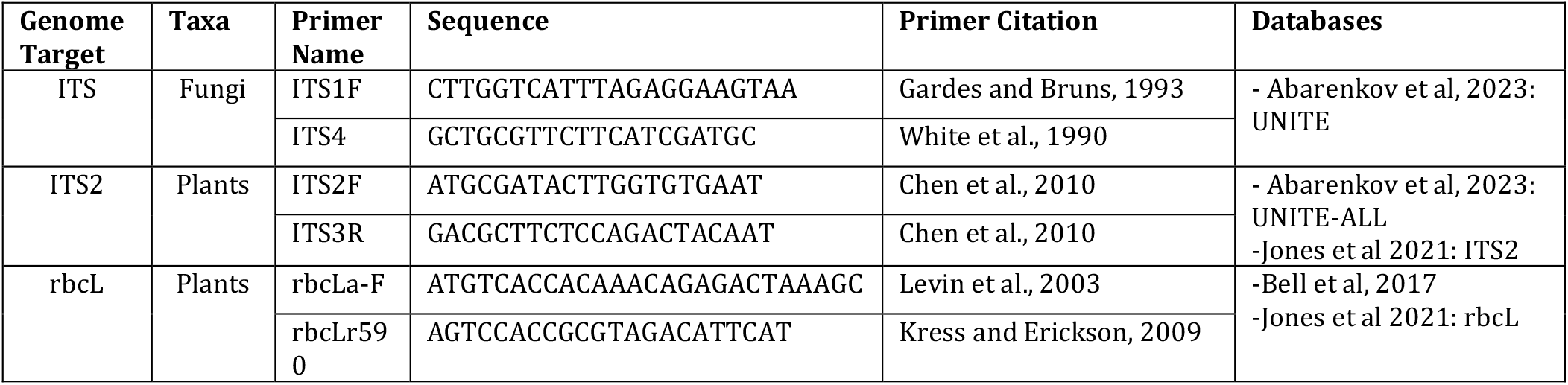
Primer pairs and suggested databases.

#### Consensus Calling

Forward and reverse chromatograms are loaded and quality trimmed using PHRED quality score (Ewing et al. 1998; Ewing and Green 1998) and chromatogram peak heights. A sliding window approach is used to remove low-quality regions. High-quality reads are aligned using Biopython’s PairwiseAligner() to produce consensus sequences. If only one read passes quality filters, that one alone is used; if none of them pass the quality filters, then the sample is discarded. This ensures high confidence in downstream classification.

#### Classification

Consensus or individual reads are classified taxonomically using blastn (Altschul et al. 1990). Blast (Basic Local Alignment Search Tool) hits are filtered based on length, percent identity, and coverage thresholds. A taxonomic group is assigned by finding agreement across top hits within a differential of 0.5% (by default) of the highest percent identity. The output taxonomic rank is truncated at the last common level shared by all top hits and further constrained by minimum percent identity cutoffs (e.g., 97% for species-level). Plant and fungal ITS sequences classified with the UNITE database (Abarenkov et al. 2023) to the species level are assigned an SH number or a genus-species binomial, if using a database other than UNITE. Fungal samples that do not achieve the 97% similarity species-level classification or which have not been assigned a species-level taxonomy are passed to the clustering phase for further resolution.

#### Clustering

Samples passed to the clustering workflow are quality-filtered and then processed through ITSx (Bengtsson-Palme et al. 2013) to retain only those with full ITS regions. These are then clustered into operational taxonomic units (OTUs) using vsearch (Rognes et al. 2016) at a default identity threshold of 97%. Each cluster is assigned a name in the format “denovo_X”, representing novel taxa not present in the reference database. These clusters, along with species-level hits from the classification step, constitute the full set of species in each study/project.

#### Salvaging

The salvaging workflow attempts to recover sequences that failed initial classification or clustering due to sequencing errors and low sequence quality. This is achieved by ‘blasting’ the as-yet unclassified samples against the reference sequences that have already be positively identified in the study and tolerating low percent identify of matches (assuming many sequencing errors occur in these samples). In order to help prevent artificially inflated diversity estimates, the user can define “salvage groups” that restrict the possible classifications that could be assigned to one of those already present among sample within the same group (often a soil core, sampling locality, or geographic area). This minimizes misclassification due to low-quality data and prevents inflation of local diversity estimates.

#### Functional Annotation

Fungal sequences with taxonomic assignments at a certain taxonomic level (e.g. mostly species or genus) can be functionally annotated using FUNGuild (Nguyen et al. 2016). This optional step provides ecological context such as trophic mode or lifestyle, aiding downstream biodiversity analysis.

#### Output

The primary output is a classification table detailing consensus results, taxonomies, clusters, and functional annotations. A log file summarizes metrics describing how many samples were successfully classified, clustered, salvaged, and functionally annotated. The output package also includes FASTA/FASTQ files, BLAST results, clustering output, and FUNGuild results (for fungi) for transparency and manual validation.

### Technical Specification

#### Programming Language

Python3.11

#### Operational system

Linux/Unix

#### Dependencies

BioPython* (1.83): https://biopython.org/wiki/Getting_Started

blastn* (2.14.0): https://blast.ncbi.nlm.nih.gov/doc/blast-help/downloadblastdata.html

ITSx* (1.1.3): https://microbiology.se/software/itsx/

vsearch* (2.21.1): https://github.com/torognes/vsearch

FUNGuild (1.1.0): https://github.com/UMNFuN/FUNGuild

*included in a conda environment called MycorhizaTracerEnv. Accessed with environment.yaml.

#### Input files

Metadata sheet specifies the sample name and the associated DNA sequencing files (chromatograms) as well as the groups for salvaging and other relevant information as shown in Table 2.

**Table 2:** Metadata file column descriptions.

| Column Name | Notes |
| --- | --- |
| Sample_ID | A unique sample identifier |
| salvageGroup | A grouping factor such that samples in need of salvaging are restricted to taxa that already exist in the dataset. Keeps samples with poor sequencing quality from adding diversity to sites. |
| directory | The directory in which that the raw reads are stored.<br>The full filepath for the sequencing file will be specified as:<br><--prefix>/<directory>/<XXX_For_file><br>Both --prefix and directory can be empty if the entire filepath is stored in <XXX_For_file>. |
| ITS_For_file | The forward ab1 file for the fungal full ITS fragment |
| ITS_Rev_file | The reverse ab1 file for the fungal full ITS fragment |
| ITS2_For_file | The forward ab1 file for the plant ITS2 fragment |
| ITS2_Rev_file | The reverse ab1 file for the plant ITS2 fragment |
| rbcL_For_file | The forward ab1 file for the plant *rbcL* fragment |
| rbcL_Rev_file | The reverse ab1 file for the plant *rbcL* fragment |

**Table 3:**
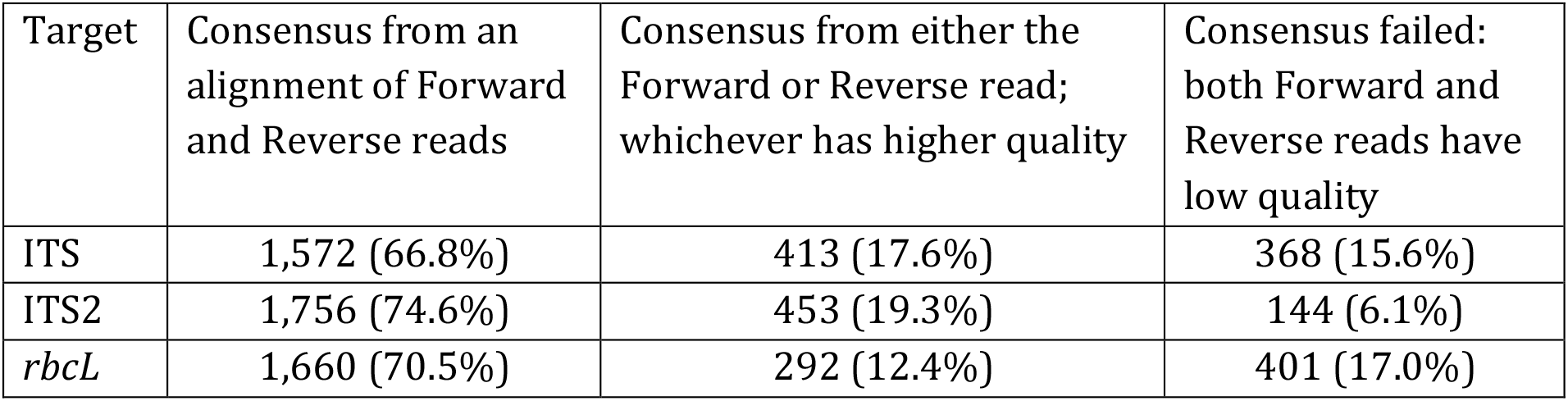
Consensus generation statistics.

DNA sequence chromatograms in ab1 format.

#### Suggested Databases

For fungal ITS: UNITE found at https://doi.plutof.ut.ee/doi/10.15156/BIO/2959333 (Abarenkov et al. 2023)

For plant ITS2: ITS2 sequences described in (Jones et al. 2021) and deposited in BOLD (Ratnasingham et al. 2024) and NCBI or UNITE-ALL found at https://doi.org/10.15156/BIO/2959334 (Abarenkov et al. 2023)

For plant rbcL: rbcL sequences described in (Jones et al. 2021) and deposited in

BOLD (Ratnasingham et al. 2024) and NCBI or rbcL sequences described in (Bell et al. 2017) and found at: https://figshare.com/articles/dataset/rbcL_July_2021/14936007?backTo=/collections/rbcL_reference_library_July_2021/5504193

### Repository

#### Type

Git

#### URL

https://github.com/MycologyKew/MycorrhizaTracer

### Usage licence

MIT license

### Implementation

MycorrhizaTracer is implemented as an executable Python script (“Classify_Sanger_OTUs.py”) with a command-line interface. The pipeline is distributed with a conda environment containing all necessary packages and executables, with the one exception of FUNGuild (Nguyen et al. 2016), ensuring reproducible setup and deployment. Each major component of the workflow—input/output handling, consensus generation, taxonomic classification, clustering, salvaging, and functional annotation—is configurable via command-line arguments, allowing the user to fine-tune pipeline behaviour to suit specific datasets or experimental goals.

#### Metadata and databases

A metadata file is required to specify the location of the chromatogram files and define salvaging groups. These groups are used during the optional salvaging step to limit taxonomic inference from low-quality sequences to predefined ecological or geographic contexts. The expected metadata format is described in Table 2.

While any *.fasta* file with sequences which are targeted by the primers chosen will work as a reference database, the header lines of each sequence have a specific required format. The databases for fungal ITS (*--ITS_db*), plant ITS2 *(--ITS2_db*), and plant rbcL (*--rbcL_db*) must have headers formatted as found in the UNITE general release *.fasta* (Abarenkov et al. 2023), though some of the fields are never used in the pipeline here:

> “*Genus_species|{unused_field}|{SH_number_or_similar}|{unused_field}|k_kingdom;p_phylum;c_class;o_order;f_family;g_Genus;s_Genus_species”*.

#### Consensus Generation

The pipeline begins by loading forward and reverse chromatograms and applying quality trimming. Trimming is based on two metrics: PHRED quality scores and chromatogram peak height. These are evaluated using a sliding window approach, with user-definable parameters such as --qual (minimum PHRED score), --window_size (window length), and --stddev_cutoff (chromatogram peak height threshold, in standard deviations below the mean). The window is moved base by base across the sequence ends, trimming low-quality regions until both metrics are satisfied.

After trimming, reads are assessed against a minimum length requirement, set via the --min_read_length flag. If both forward and reverse reads exceed this threshold, they are aligned using a local pairwise aligner from Biopython’s Bio.Align module. A consensus sequence is then generated: matching bases are retained with the higher quality score, while mismatches are replaced with an “N”. If only one read passes filtering, it is used directly as the representative sequence for the sample and will here be referred to as a “consensus”. Samples with no passing reads are omitted from downstream analyses.

#### Taxonomic Classification

Taxonomic classification is performed using blastn (Camacho et al. 2009), with the consensus sequence or high-quality individual reads queried against a user-specified database. To refine the classification to regionally relevant taxa, the user may optionally supply species lists (via --species_list_ITS, --species_list_ITS2, or --species_list_rbcL). These lists must contain genus and species columns, and only database entries with exact matches are retained during classification.

BLAST hits are initially filtered by alignment length (--minMatchLength, default 50bp) and coverage thresholds for both query and subject sequences (--minQCOV and --minSCOV). A match passes if either coverage exceeds 0.95 or both exceed the defined thresholds. Additional filtering steps are available: the --no_Incertae_Sedis flag removes blast hits with ambiguous taxonomy from consideration (unless every hit is thus removed, in which case the original list of hits is used), while --adjustPidents adjusts identity scores to refine the percent identity of partial alignments.

The highest percent identity from the remaining hits is saved and any hits within a user-defined identity differential (--pidentDiff, default 0.5%) of this top percent identity are selected as candidate matches. These are compared taxonomically from kingdom to species. Where disagreement is found, the taxonomic ranking is truncated to the last level at which all hits agree. Final classification is further constrained by a series of minimum identity cutoffs for each rank (e.g., --min_pident_species, default 0.97). The final classification is stored for each sample, and the top BLAST hit’s full taxonomy is also recorded separately for users requiring higher resolution who can tolerate lower accuracy, though this should be interpreted with caution.

Plant and fungal sequences that receive a species-level match against the UNITE database are assigned a UNITE SH number. For plant sequences classified using the *rbcL* region, a genus-species label is used instead. Fungal samples failing to achieve species-level resolution are passed to the clustering phase.

#### Clustering

Sequences not classified to the species level at the designated cut off (the default of 97.0 was used in this case study) enter the clustering workflow. These are subjected to stringent quality and length filtering using thresholds set by --minSeqLengthForCluster (default 300 bp) and --minAverageQualityForCluster (default PHRED 20). High-quality reads are then processed with ITSx to extract the ITS region (ITS1, 5.8S, ITS2); sequences lacking a full ITS region are saved for later salvaging (optional).

Valid sequences are clustered using vsearch --cluster_fast with an identity threshold defined by --percentIdentityForCluster (default 97.0). The resulting OTUs are given names of the form denovo_X, representing species not found in the original reference database. These clusters are comparable to UNITE’s SH numbers and serve as *de novo* taxonomic units for diversity analyses.

#### Salvaging

The optional salvaging workflow, triggered via the --salvage flag, attempts to recover information from sequences excluded during consensus or clustering. These sequences are aligned against a custom BLAST database composed of SH reference sequences and cluster centroids identified earlier in the pipeline. Each salvaged sample is assigned the taxonomy of its best hit by e-value, provided that a sample from the same salvage group has already been assigned that taxonomy. This requirement ensures that poor-quality reads do not introduce artificial taxa to the dataset and helps maintain local consistency in diversity metrics.

#### Functional Annotation

The final step, enabled with --run_funguild, performs functional annotation of fungal taxa using FUNGuild (Nguyen et al. 2016). This tool assigns trophic mode, lifestyle, and ecological function based on the consensus taxonomy. The parsed guild assignments and confidence scores are summarized in the output, with the full FUNGuild report also included for reference.

## Results and Conclusion

### Overview of sample dataset

The dataset used for development and testing of this pipeline contains 2,357 individual ectomycorrhizas collected from 63 soil plots across England, for fungal and plant identification following previously published methods (Suz et al. 2014; van der Linde et al. 2018). Soil samples were collected through Defra’s Natural Capital and Ecosystem Assessment (NCEA) programme. The PCR amplification used the fungal specific primer ITS1F (Gardes and Bruns 1993) and the eukaryotic primer ITS4 (White et al. 1990), and two sets of primers for plants: ITS2F and ITS3R (Chen et al. 2010) targeting ITS2 and rbcLa-F (Levin et al. 2003) and rbcLa590 (Kress et al. 2009) which target *rbcL*. The PCR products were visualised on an agarose gel and submitted for bidirectional Sanger sequencing to Azenta UK Ltd (GENEWIZ Sanger Sequencing Laboratory, Takeley, United Kingdom). The resulting 14,134 (a few samples were not sequenced at one of the redundant plant targets) chromatograms contain a range of sequence lengths and qualities as may be expected from environmental samples (Fig. 2). We have included poor-quality sequences here to showcase the pipelines’ ability to find and appropriately handle such occurrences. The anonymized data and two metadata sheets are provided in the supplement: one with site-specific salvage groups (63 groups) and one with country-specific salvage groups (1 group). For the statistics reported below, version 2.2.1 was run once with each metadata sheet using the flags --metadata, --prefix, --dirDepth, --salvage, --run_funguild, and --sequence_stats. The databases used defined with --ITS_db, --ITS2_db, --RBCL_db, are those referenced above.

**Figure 2:**
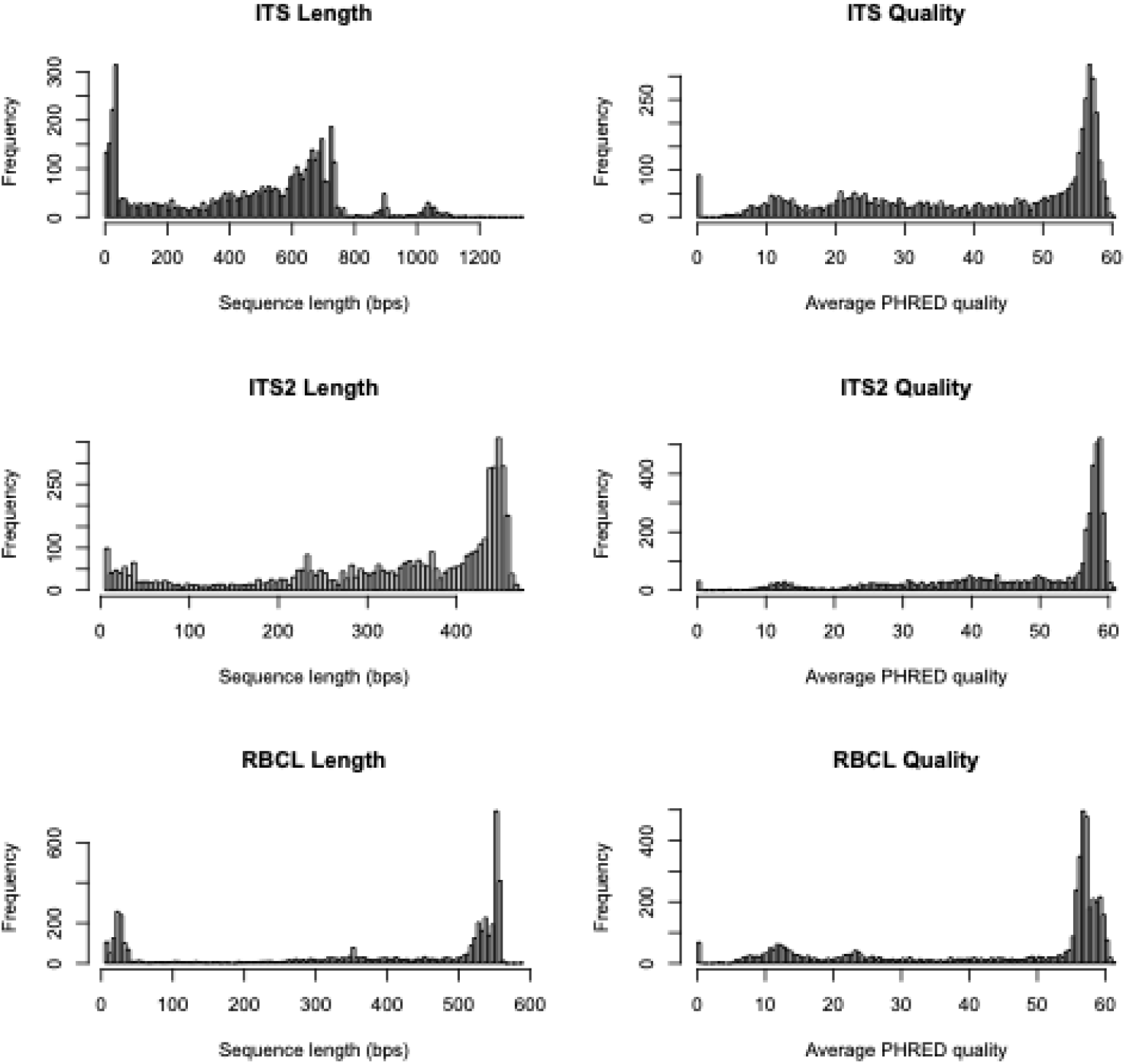
Post-filtering sequence statistics for test samples (full ITS for fungi, ITS2 and *rbcL* for plants). These samples span a range of qualities and lengths and include many of the variants that may appear in chromatograms, such as high quality suddenly dropping due to an indel in the sample, and samples whose sequencing failed all together.

### Consensus Generation Efficiency

Of the 2,357 samples across all three target regions, consensus sequences were successfully generated for 88% of forward-reverse sequence pairs (86% for fungal full ITS, 94% for plant ITS2, and 83% for *rbcL*). The majority (85%) of successful consensus sequences were constructed from both forward and reverse DNA sequences, while the remaining 15% were based on a single sequence (either forward or reverse). These numbers are primarily dependent on the sample and sequencing quality and successful consensus building is unsurprisingly far more likely with high-quality sequences.

### Taxonomic classification results

In order for a sample to be classified, it had to exceed the user-defined match length (default = 50 bp) and coverage thresholds (default 0.3 for scov and qcov), match the database at or above the user-specified percent identity threshold (default 97% for species and 94% for genus) and all blast hits within a percent identity differential (default = 0.5%) of the highest percent identity must have had the same taxonomy. For each target region, ca. 17% of samples (19.6% ITS, 15.4% ITS2, 17.7% *rbcL*) were not classified to any taxonomic rank due to poor sequencing quality. Of those that had some classification with the initial blast, 62% (fungal I TS), 54% (plant ITS2) and 47% (plant *rbcL*) achieved classification to the species level and 79% (ITS), 65% (ITS2) and 73% (*rbcL*) were classified at the genus level (Fig. 3). These taxonomic classifications are strongly dependent on the database completeness (i.e.: the proportion of the species that exist which have records in the database).

**Figure 3:**
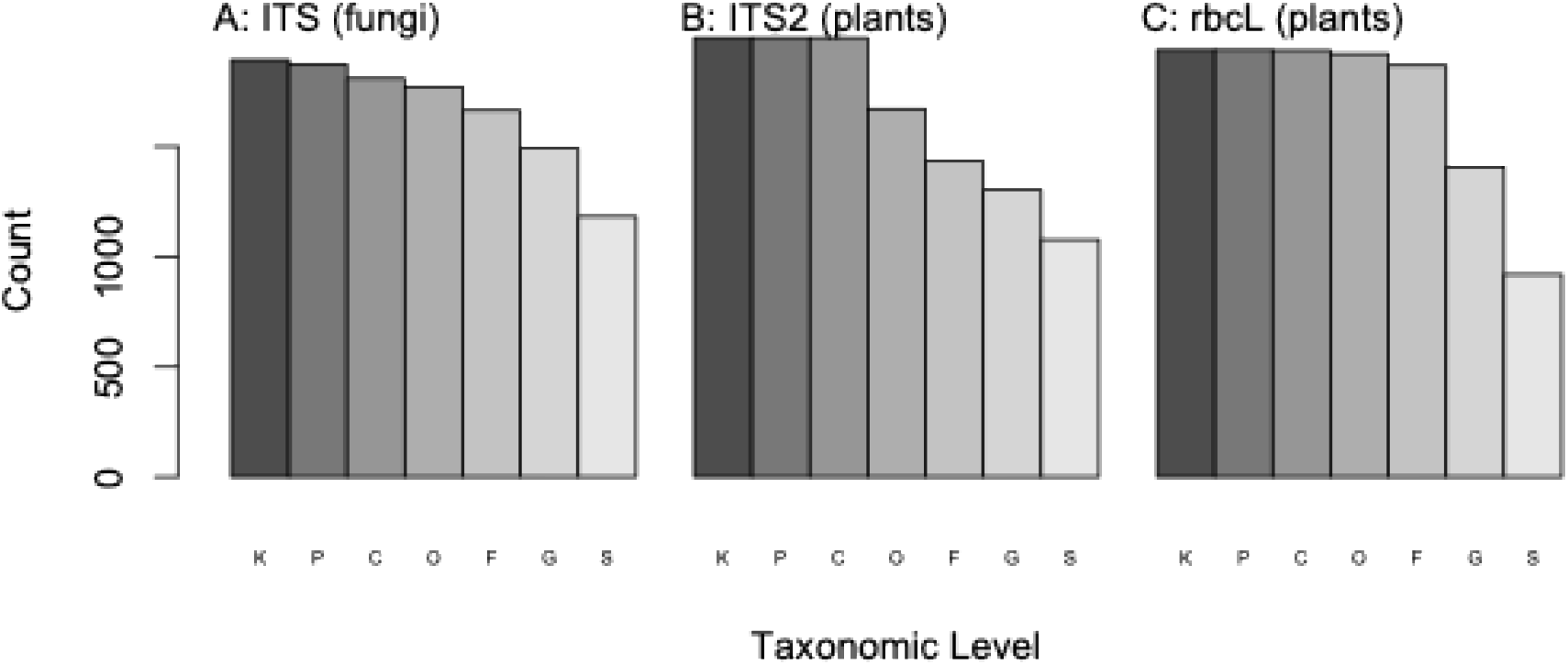
Taxonomic classification success. (A) Fungal ITS with UNITE, (B) plant ITS2 with BOLD, and (C) *rbcL* with BOLD. Default percent identity thresholds were used: <u>K</u>ingdom:>80%, <u>P</u>hylum:>80%, <u>C</u>lass:>80%, <u>O</u>rder:>85%, <u>F</u>amily:>90%, <u>G</u>enus:>94%, <u>S</u>pecies:>97%.

### Operational Taxonomic Units

Operational taxonomic units were defined as the combination of (1) the species names of samples with species-level classifications (or SH numbers) and (2) the cluster centroids. For fungi, OTUs are a combination of species-level classifications from the blastn (i.e.: SH numbers) and the results of clustering the remaining high-quality sequences at 97% identity. The clustering step is constitutive for fungi, as many fungal species are yet undescribed and do not appear in the databases, but is optional for plants and left to the users’ discretion. The pipeline identified 677 fungal OTUs (samples = 1,796, mean size = 2.6, max size = 117, singletons = 467), 19 plant OTUs based on ITS2 (samples = 1,077, mean size = 56.7, max size = 753, singletons = 8) and 15 plant OTUs based on *rbcL* (samples = 919, mean size = 61.2, max size = 637, singletons = 5).

### Salvaging performance

Our salvaging workflow attempts to rescue low-quality sequences by using a blastn search to existing OTUs with a lenient use of the highest e-value to decide the best hit and then a stringent filter for whether another sample in the same salvage group has already been assigned to that OTU. The highest e-value is used to allow tolerance for poor-quality sequences that may contain sequencing errors or ambiguities. The strict already-present filter is then used to keep low-quality sequences from increasing the alpha diversity of the site; low-quality sequences should not be the only instance of an OTU at a site. The number of successful salvages is thus mainly constrained by two factors: (1) the sequence quality impacting blast’s ability to identify a potential hit and (2) the size of the dataset at each site (i.e., the number and size of salvaging groups specified in the metadata). Having a finer-scale grouping structure (i.e. using study sites as groups) may result in fewer successful salvages than a coarser-scale grouping structure (i.e. using countries as groups). When using individual sites for salvage groups (n=63 sites), 59 of the attempted 256 fungal samples were salvaged, whereas when using the country as a group (n=1 country; all sites are in the UK) then 121 of 256 samples were successfully salvaged. The decision as to how fine to set the salvage grouping is an important consideration for those using the pipeline, as it will influence the balance between treating rare (or singleton) sequences as novel taxa or discarding them as a low-quality sequencing attempts. Indeed, singleton sequences in any ecological dataset are always worth special consideration, as they are likely to be enriched for sequencing errors, but rare and novel species (particularly for microbes) exist and removing all singletons may result in underestimated diversity. In our study dataset each sample is an individual ectomycorrhiza whose morphology was confirmed by microscopy, so high quality singletons assigned as ectomycorrhizal fungi are not removed from the dataset.

### Comparison with other existing pipelines

Sanger sequencing has been used for decades and a few pipelines do exist that are designed with a focus on Sanger data. Table 4 presents a comparison between MycorrhizaTracer and other pipelines with similar functionality. Most similar is SangerFlow (Prodhan et al. 2025), a nextflow-based program designed and optimised for high-throughput pest and pathogen screening which differs from MycorrhizaTracer in that it does not cluster the samples and does no functional annotation of fungal samples. Also similar to MycorrhizaTracer is SCATA (Durling et al. 2011) which is also optimised for fungal ITS barcoding. SCATA is only available as a web-interface and so may be more accessible in the absence of bioinformatic expertise, but it does not accept chromatograms necessitating the major bioinformatic hurdle of sequence editing and consensus building. Neither does SCATA interface with FUNGuild for functional annotation. PipeBar (Oliveira et al. 2018) and SangerAnalyzeR (Chao et al. 2021) are both designed for processing Sanger chromatograms at scale, but both stop at the consensus generation stage, and thus classification and clustering are left to the user. To our knowledge, MycorrhizaTracer is the only pipeline that takes raw chromatograms through to the functional annotation stage.

**Table 4:**
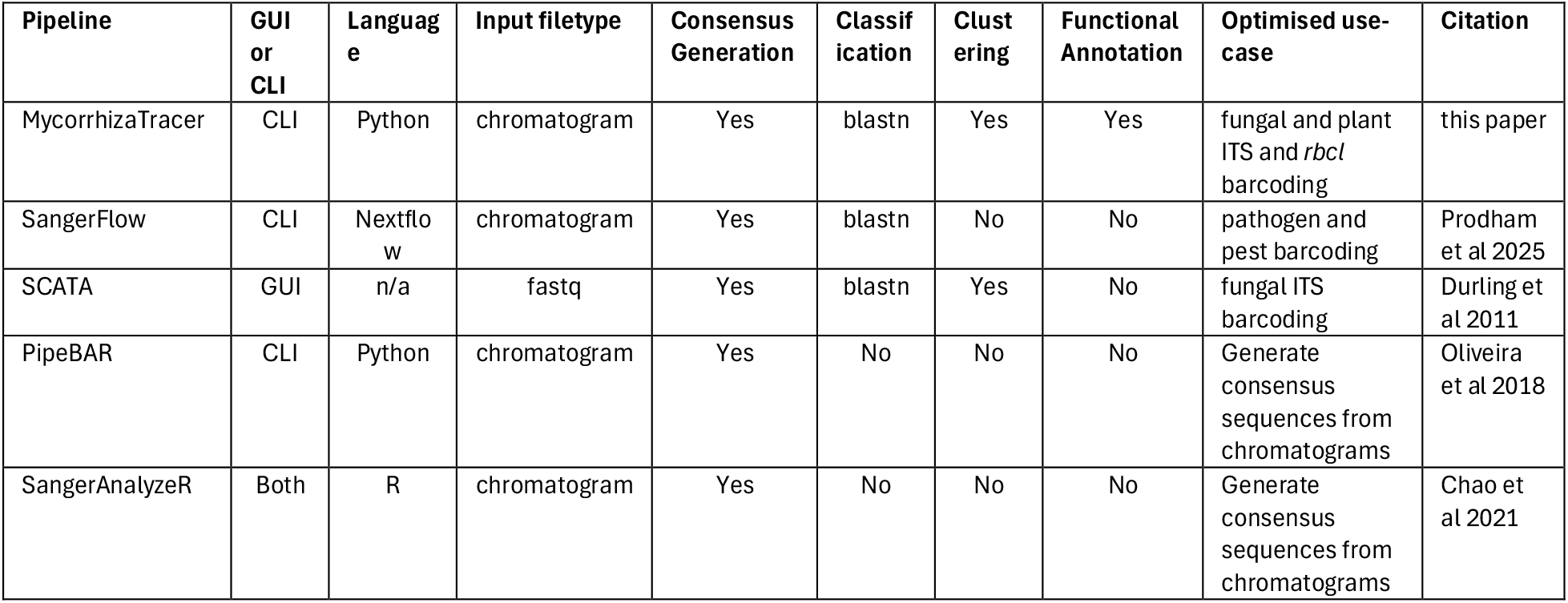
Similar programs developed to process Sanger DNA sequences.

### Limitations and Considerations

Despite its strengths, MycorrhizaTracer has some limitations that warrant consideration. Most notably, many of the failures to assemble a consensus or achieve species-level classification stem from poor-quality input sequences. This underscores the importance of generating high-quality chromatograms at the sequencing stage, as downstream performance is tightly coupled to input data quality. In addition, taxonomic classification accuracy remains limited by the breadth and curation of underlying reference databases. Unknown or misannotated taxa can lead to ambiguous or incorrect matches. Furthermore, the pipeline’s command-line interface, while flexible, may be less accessible to users without prior bioinformatics experience. The salvage workflow, though designed to prevent overinflated diversity estimates, also depends on the completeness and correctness of sample metadata for salvage grouping. Finally, functional annotation is limited to fungal taxa interpretable by FUNGuild, and confidence in guild assignment can vary widely depending on taxonomic resolution and coverage, thus expert opinion, visual inspection of the samples, a thorough understanding of the literature, and a detailed understanding of the focal ecology are all important additions to this and any pipeline.

### Implications and future directions

MycorrhizaTracer is an efficient and scalable tool for processing and classifying large numbers of plant and fungal DNA chromatograms. It is fully automated, taking raw chromatograms through to functionally annotated classified taxa, while reducing the need for manual inspection of trace files. This allows for consistent, unbiased processing across large datasets. All critical thresholds (quality, coverage, identity cutoffs, taxonomic filters) are exposed as command-line arguments, allowing users to tune the pipeline for different barcoding regions, target taxa, and sample quality levels without modifying the code. The optional inclusion of user-supplied species lists for regional taxa ensures that classification results are ecologically plausible and aligned with expert expectations. This is particularly useful for localized biodiversity assessments. The (Nguyen et al. 2016), allowing ecological interpretation of the taxonomic output for fungi. This helps bridge taxonomy and function, streamlining downstream ecological analyses when sequences can be assigned taxonomy at least at genus level. The pipeline generates detailed output files, including intermediate FASTA/FASTQ, raw BLAST results, clustering summaries, and full FUNGuild outputs. This transparency supports reproducibility and provides users with checkpoints for quality assurance or custom downstream workflows. In benchmark tests, 2,357 samples (14,134 chromatograms) were processed in under 20 minutes on a consumer-grade laptop (MacBook Air, 6 CPUs, 16 GB RAM, Apple M3), demonstrating high efficiency, even without access to a high-performance computer.

## Acknowledgements

This manuscript is led by The Mycorrhiza Lab at RBG Kew and funded by the UK Government through Defra’s Natural Capital and Ecosystem Assessment programme. We would like to thank the lab team who processed the samples that we used for a test-data set, including: R. Jarvis, O. Lindsay, G. Meadows, M. Rickards, E. Serpell, and B. Underwood.

The Natural Capital and Ecosystem Assessment (NCEA) programme is Defra’s largest Research and Development (R&D) programme and the most ambitious assessment of the state of nature ever undertaken in the UK. NCEA is assessing England’s land, freshwater, and coastal ecosystems to produce a baseline of our natural assets by 2029, enabling a natural capital approach to policy and investment.

## Author Contributions

**T.D.B.** developed the code and wrote the manuscript.

**T.W., R.B., I.T.**, and **R.Go.** beta-tested, error-checked, and troubleshot the code.

**R.Ga., G.D. and L.M.S**. contributed to planning the codes’ organisational structure and development strategy.

**C.A., J.K., M.B.**, and **L.M.S.** provided subject-matter expertise and valuable input on the codes’ structure and function.

